# Wearable Estimation of Knee Joint Contact Forces: SiMPLeSENSE, a Simulated-IMU Pipeline Validated Against *In Vivo* Loads

**DOI:** 10.64898/2026.11.03.763565

**Authors:** Alessandro Castellaz, Sabrina Hörmann, Ajay Seth, Bert-Jan F. van Beijnum, Frank J. Wouda

**Affiliations:** University of Twente, Drienerlolaan 5, 7522 NB, Enschede, Netherlands; Delft University of Technology, Mekelweg 5, 2628 CD, Delft, Netherlands

**Author notes:** Corresponding author. *Email address:* (Alessandro Castellaz). These authors contributed equally to this work (Alessandro Castellaz, Sabrina Hörmann).

**Keywords:** Knee contact force, Inertial measurement unit, Musculoskeletal modeling, Wearable sensors, Gait

## Abstract

Knee joint compressive forces (KCF) are central to understanding joint loading and degeneration, but they can be measured directly only with instrumented implants. Musculoskeletal models provide a non-invasive alternative, and inertial measurement units (IMUs) could make such estimates portable. How-ever, no dataset combines IMU signals with *in vivo* implant forces, so the accuracy of IMU-based KCF remains unverified. We developed SiMPLeSENSE, a framework that derives virtual IMU orientations from optical motion capture using subject-specific SMPL meshes. Using the CAMS-Knee dataset, we reconstructed kinematics with OpenSense and estimated KCF in OpenSim with a Rapid Muscle Redundancy solver, comparing the estimates with *in vivo* measurements during level walking and stair descent. We also assessed robustness to sensor placement error by adding empirically derived thigh and shank orientation noise (for two cases). Across subjects and activities, the simulated-IMU estimates closely tracked the optical mocap-driven estimates and reproduced the *in vivo* force pattern, with the agreement against the optical reference being higher for stair descent. Overall, the estimated forces matched the *in vivo* measurements with a median RMSE of about 0.3 to 0.5 BW and correlations near 0.95. Adding realistic orientation error increased RMSE only modestly, whereas doubling it markedly overestimated peak force, most evident for stair descent. These results indicate that the approach is robust to substantial changes in input kinematics, supporting a laboratory-independent setup combining inertial motion capture, a small EMG set, and instrumented force shoes for home and clinical monitoring of knee loading.

## Introduction

Excessive or unevenly distributed knee joint loading is associated with cartilage damage and osteoarthritis progression (Kumar et al., 2013; Andriacchi & Mündermann, 2006). Knee joint compressive forces (KCF), vary substantially across daily activities. *in vivo* measurements show that those forces reaches roughly two to three times body weight during level walking and exceeds three times body weight during stair climbing (Kutzner et al., 2010; Heinlein et al., 2009). Therefore, reliable KCF estimation is of considerable clinical relevance to understand how loading changes with activity and retraining, which can be effective treatments for OA (Uhlrich et al., 2025).

Direct measurement of KCF requires an instrumented implant. The CAMS-Knee dataset provides such *in vivo* forces (Taylor et al., 2017), and datasets of this kind are the gold standard for validating musculoskeletal models (Fregly et al., 2012; Heinlein et al., 2009). Because the implants are invasive, their use is limited to a small number of participants who have undergone total knee arthroplasty, which constrains broader methodological validation.

A widely used non-invasive alternative estimates joint forces with muscu-loskeletal models driven by optical motion capture and force plates. Platforms such as OpenSim combine measured kinematics with a biomechanical model to compute joint moments and contact forces (Delp et al., 2007; Seth et al., 2018), with estimates that agree reasonably (RMSE ≈ 0.4 BW, *ρ* ≈ 0.9) with implant measurements during walking and other functional tasks (Imani Nejad et al., 2020; Pelegrinelli et al., 2023). A limitation of optimization-based solvers is that, by minimizing muscle effort, they neglect co-contraction, the simultaneous activation of agonist and antagonist muscles, which adds to knee loading (Belli et al., 2023; Rudolph et al., 2000). Incorporating measured EMG addresses this, and EMG-driven and EMG-informed models have repeatedly improved KCF estimates, particularly for atypical gait or high co-contraction (Lloyd & Besier, 2003; Manal & Buchanan, 2013; Sartori et al., 2012; Pizzolato et al., 2015; Hoang et al., 2019; Princelle et al., 2025). Following this approach, Hörmann et al. (2026) added a co-contraction index (CCI) term to the muscle redundancy solver, constraining the model toward the co-contraction measured between vastus lateralis and gastrocnemius medialis, and reported improved KCF estimates.

Optical systems, however, require a dedicated laboratory, fixed cameras, and trained personnel, which restricts their use outside controlled settings. Wearable inertial measurement units (IMUs) offer a portable, lower-cost alternative, and IMU-based kinematics agree well with optical motion capture for lowerlimb joint angles across many activities (Picerno, 2017; Karatsidis et al., 2017; Schepers et al., 2018). Several studies have used IMU-derived kinematics in musculoskeletal models to estimate knee moments and contact forces. Konrath et al. (2019) reported good agreement between IMU-and laboratory-based KCF during daily activities, and Di Raimondo et al. (2023) found comparable peak forces from the two pipelines. All of these studies, however, validate against laboratory-derived estimates rather than *in vivo* implant forces, so the absolute accuracy of IMU-based KCF remains uncertain (Imani Nejad et al., 2020; Konrath et al., 2019).

A central obstacle is the lack of datasets that record IMU signals and *in vivo* forces on the same trials. Simulating IMU data from optical motion capture provides a controlled solution, retaining the *in vivo* ground truth while allowing an inertial pipeline to be evaluated under known conditions. CROMOSim applies this principle, generating virtual inertial signals from motion data (Hao et al., 2024), and simulated IMU data have also been used to train and test gait pipelines when real recordings are limited (Dorschky et al., 2020; Mundt et al., 2020). Realistic virtual sensors are enabled by statistical body models, in particular the Skinned Multi-Person Linear (SMPL) model and its dynamic soft-tissue (DMPL) extension, which reconstruct the body surface from marker data so that sensors can be placed on it (Loper et al., 2015; Mahmood et al., 2019).

In this study we adapt the CCI-informed pipeline of Hörmann et al. (2026) by replacing its optical input with simulated IMU data while retaining measured GRFs, to assess the effect of IMU orientation accuracy on KCF estimates. Using optical data from the CAMS-Knee dataset, we reconstruct each subject with SMPL, generate virtual IMU orientations, reconstruct full-body kinematics with OpenSense (Al Borno et al., 2022), and compute KCF in OpenSim using the CCI-informed and minimum activation RMR solver. The resulting forces are compared directly with the *in vivo* implant measurements across several activities of daily living. We then evaluate the robustness of the method to sensor placement error by adding realistic orientation noise to the thigh and shank virtual IMUs (Fan et al., 2022), including an extreme case in which the noise is doubled.

## 2. Methods

### 2.1 Dataset

We used the CAMS-Knee dataset (Taylor et al., 2017), which provides synchronized *in vivo* joint loading alongside optical motion capture, ground reaction forces (GRF), and surface electromyography (EMG) from six participants who had undergone total knee arthroplasty with a cemented INNEX tibial implant (Zimmer, Switzerland; five male, one female; age 68 ± 5 years; mass 88 ± 12 kg; height 173 ± 4 cm).

Whole-body kinematics were recorded at 100 Hz with a 26-camera optical system and a 75-marker set (Vicon, UK). GRF were sampled at 2000 Hz from six floor-embedded force plates (Kistler, Switzerland) and low-pass filtered at 12 Hz (fourth-order zero-phase Butterworth). Bilateral surface EMG was recorded from eight lower-limb muscles per leg (rectus femoris, vastus medialis, vastus lateralis, tibialis anterior, semitendinosus, biceps femoris long head, and the medial and lateral gastrocnemii) with a 16-channel wireless system (Trigno, Delsys, USA). Following ISEK recommendations (Merletti, 1999), the raw signals were high-pass filtered at 100 Hz, full-wave rectified, and low-pass filtered at 4 Hz to obtain linear envelopes, and the 48 ms acquisition delay was corrected.

### 2.2 Virtual IMU Pipeline (SiMPLeSENSE)

We developed SiMPLeSENSE, a framework that generates virtual IMU orientations and joint kinematics from optical motion capture data by combining parametric body modeling, mesh-driven virtual sensors, and IMU-based inverse kinematics. Because the virtual sensors come from the same trials for which *in vivo* forces exist, the pipeline can be evaluated against an *in vivo* reference under fully controlled conditions, without physical sensor hardware. This follows earlier work on simulated inertial signals for data augmentation and pipeline evaluation (Hao et al., 2024; Dorschky et al., 2019, 2020). The workflow is summarized in Fig. 1.

**Figure 1:**
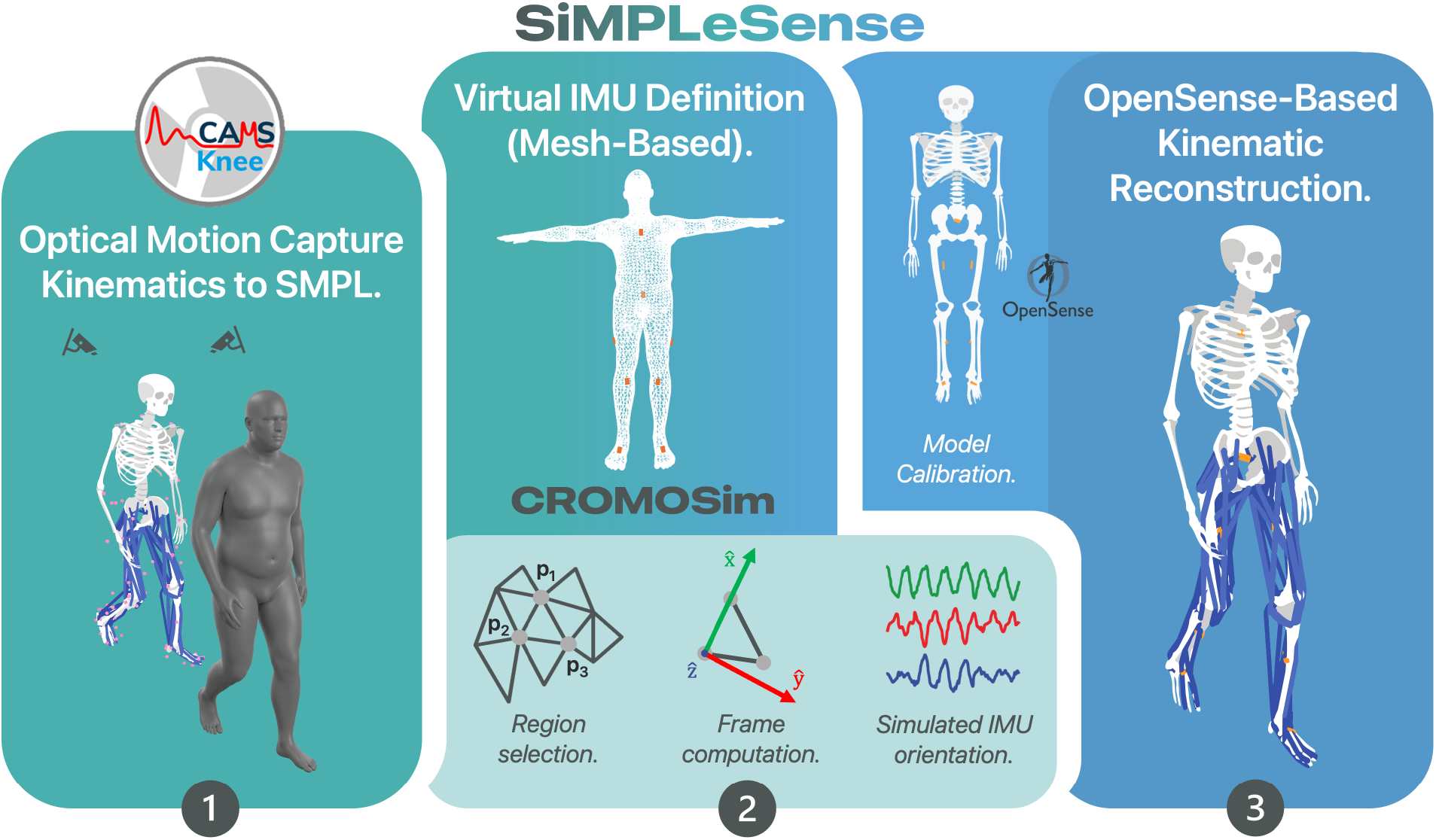
SiMPLeSENSE pipeline. Experimental motion capture data are used to drive subject-specific SMPL body models (1). Virtual IMUs are derived from mesh-based coordinate frame estimation (2). Simulated orientations are processed through OpenSense to reconstruct full-body joint kinematics (3).

#### 2.2.1 Synthetic motion generation and subject-specific modeling

In the first stage (Fig. 1, step 1), the optical motion capture data are mapped onto a subject-specific body model. We used the SMPL parametric model (Loper et al., 2015), which represents the body as a triangulated surface mesh driven by pose (***θ***), shape (***β***), and dynamic soft-tissue (DMPL) parameters. Subject-specific shape coefficients were estimated from each participant’s height, mass, and sex, using 16-dimensional shape and DMPL spaces (Loper et al., 2015; Mahmood et al., 2019). Joint kinematics from OpenSim inverse kinematics (.mot) were mapped to SMPL pose parameters through a predefined joint-axis correspondence, while global pelvis translations were omitted because, as in CROMOSim (Hao et al., 2024), OpenSense uses segment orientations and corresponding joint angles rather than the global pelvis location.

#### 2.2.2 Mesh-based virtual IMUs and orientation reconstruction

From the animated mesh we defined virtual IMUs directly on the body surface (Fig. 1, step 2), following the mesh-based virtual-sensor concept of CRO-MOSim (Hao et al., 2024). We placed one virtual IMU on each of eight body segments (torso, pelvis, and the bilateral femur, tibia, and foot), in locations analogous to conventional physical sensor placement (Schepers et al., 2018). The orientation of each sensor was then derived in three steps, shown at the bottom of Fig. 1 (step 2). First, in a region-selection step, we identified a small patch of the mesh at the sensor site and tracked the time-resolved trajectories of three non-collinear vertices **p**_1_, **p**_2_, **p**_3_ ∈ R^3^ on that patch, directly defining the sensor orientation input required by OpenSense rather than simulating raw accelerometer and gyroscope signals. Second, in a frame-computation step, we built a local orthonormal frame from these vertices at each time frame:

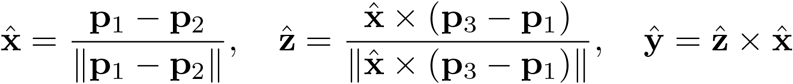

Third, the sequence of these frames over time formed the simulated IMU orientation signal. Because numerical errors and mesh noise can produce orientation matrices that are only approximately orthonormal, each matrix was projected onto the nearest valid rotation matrix using singular value decomposition (SVD). The resulting orthonormal rotation matrices were then converted to quaternions (Shuster, 1993).

#### 2.2.3 OpenSense calibration and inverse kinematics

In the final stage (Fig. 1, step 3), the simulated orientations (exported as OpenSim .sto files) were used to reconstruct full-body kinematics in OpenSense (Al Borno et al., 2022). The IMUPlacer tool first calibrated the full-body musculoskeletal model (Rajagopal et al., 2016; Lai et al., 2017; Uhlrich et al., 2022) to the virtual sensors from a static reference pose. The IMUInverseKinematicsTool then solved for the kinematics that best matched the simulated orientations for each trial. Because the mesh-based reconstruction captures soft-tissue motion but not sensor-to-segment misalignment, we added realistic orientation noise in a sensitivity analysis (Section 2.5) to test the robustness of the KCF estimates.

### 2.2 Kinetic Adjustment for Fixed-Pelvis Kinematics

Because OpenSense does not estimate the global position of the pelvis and only computes the coordinates that minimize orientation error (without using translation information), we transformed the measured GRF into a local footbased representation, eliminating dependency on global translation and providing a consistent representation relative to the contact foot. For each trial, we ran forward kinematics on the original and simulated sequences with the scaled subject-specific model to track the pose of the contact bodies (bilateral calcanei). The centre of pressure (CoP) was mapped into the foot frame under the original kinematics and re-expressed in the local frame under the simulated kinematics:

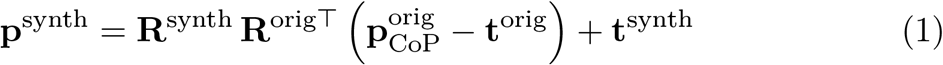

Where **R** and **t** denote the rotation matrix and position of the contact body, respectively, under the original (*orig* ) and simulated (*synth*) kinematics, 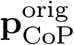 is the measured CoP position, and (·)^⊤^ denotes matrix transpose.

Ground reaction forces and free moments were rotated by only the yaw component of the relative rotation **R**_Δ_ = **R**^synth^**R**^orig⊤^, leaving the vertical components unchanged, since the gravity direction is fixed across frames.

### 2.4 CCI-Informed Muscle Redundancy Solver

We estimated KCF with the Rapid Muscle Redundancy (RMR) solver (Belli et al., 2023). The RMR solver resolves muscle redundancy by minimizing a weighted sum of squared muscle activations and reserve actuator forces:

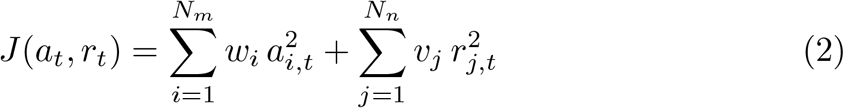

where *a*_*i,t*_ and *r*_*j,t*_ are the muscle activations and reserve actuator forces at time step *t*. We set *w*_*i*_ = 1 and *v*_*j*_ = 10 to penalize reserve actuator use (Belli et al., 2023). Unlike static optimization (Crowninshield & Brand, 1981), the RMR solver includes passive muscle fiber forces and enforces activation continuity, giving more physiologically plausible solutions (Belli et al., 2023).

More recently, Hörmann et al. (2026) extended the RMR solver with a co-contraction index (CCI) term in the cost function, constraining the optimization toward the co-contraction measured by EMG and reporting improved KCF estimates. We evaluated this CCI-informed formulation alongside the conventional *minActivation* method. The CCI-informed formulation retains the *minActivation* objective while incorporating an additional constraint based on the measured co-contraction, rather than using the EMG-driven and EMG-informed neuromusculoskeletal approaches used elsewhere to capture subject-specific co-contraction (Lloyd & Besier, 2003; Manal & Buchanan, 2013; Sartori et al., 2012; Pizzolato et al., 2015). Following Hörmann et al. (2026), the extended cost function adds a tracking term that minimizes the squared error between simulated and measured co-contraction:

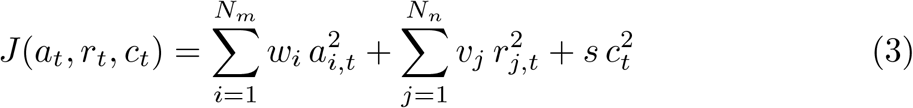

where 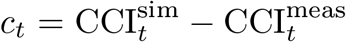 and *s* = 2 prioritizes fidelity to the measured pattern. The CCI was computed for the vastus lateralis and gastrocnemius medialis antagonistic pair, identified by Hörmann et al. (2026) as showing the largest discrepancy between measured and RMR-predicted co-contraction under minimum-activation optimization. Following Rudolph et al. (2000):

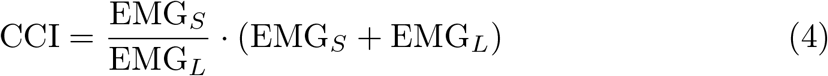

where EMG_*S*_ and EMG_*L*_ are the smaller and larger activations of the pair, which constrains the CCI to the range [0, 2] and prevents division by zero. Once we computed the muscle activations, we computed the KCF through Opensim Joint Reaction Analysis tool.

### 2.5 Sensitivity Analysis and Comparison Metrics

Although SiMPLeSENSE derives orientations from mesh-based segment tracking, it does not fully reproduce the sensor-to-segment misalignment and orientation-estimation errors of physical IMUs (Hao et al., 2024; Laidig & Seel, 2023). To quantify the sensitivity of KCF to these errors, we perturbed the thigh and shank virtual IMU orientations. Because the knee angle is defined as the shank orientation relative to the thigh, errors in these two segments propagate directly and exclusively into knee kinematics and thus KCF (Fan et al., 2022).

Error magnitudes were taken from Fan et al. (2022), who reported empirical sensor-to-segment misalignment and orientation-estimation errors for the thigh and shank during dynamic tasks. The combined per-axis error (misalignment plus orientation error) is given in Table 1.

**Table 1:** Gaussian noise parameters (mean ± std, in degrees) applied to thigh and shank virtual IMU orientations, derived from Fan et al. (2022).

| Segment | Roll | Pitch | Yaw |
|---|---|---|---|
| Thigh | 3.5 ± 3.1° | 2.1 ± 1.2° | 5.0 ± 3.3° |
| Shank | 3.6 ± 2.6° | 2.1 ± 1.5° | 3.9 ± 2.8° |

For each subject and activity we drew one angle per axis from the corresponding Gaussian distribution and composed them as an intrinsic XYZ (roll– pitch–yaw) rotation (Shuster, 1993):

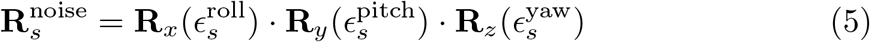

where 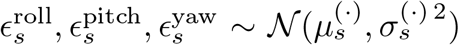 for segment *s* ∈ {thigh, shank} (Table 1). The perturbation was applied as a right-multiplication of the orientation quaternion at every frame, with the rotation magnitude sampled from the Gaussian distributions reported in Table 1, mimicking a rigid sensor-to-segment misalignment (Fan et al., 2022):

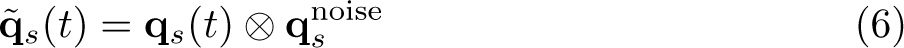

where **q**_*s*_(*t*) is the original quaternion, 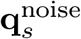 corresponds to 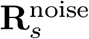, and ⊗ denotes quaternion multiplication. Only the bilateral femur and tibia IMUs were perturbed; torso, pelvis, and calcanei were left unchanged.

Calibration (IMUPlacer) used the unperturbed orientations, while the perturbed orientations entered only the inverse kinematics step. This prevents the placer from compensating for the injected noise and isolates the effect of misalignment on downstream KCF (Al Borno et al., 2022). The perturbed kinematics were then passed through the full RMR pipeline. To probe robustness under more severe conditions, we repeated the analysis with the error parameters (mean and standard deviation) doubled, an extreme scenario beyond the magnitudes typically reported (Fan et al., 2022; Jiang et al., 2022).

We compared the nominal and perturbed estimates against the *in vivo* implant measurements (Taylor et al., 2017) using two metrics: the root mean square error (RMSE) in units of body weight (BW) for absolute agreement, and the Pearson correlation coefficient *ρ* for waveform shape. Both were computed per trial and averaged across trials and subjects, reported separately for level walking and stair descent.

## 3. Results

Figure 2 compares the reconstructed kinematics with the optical reference for the worst-and best-reconstructed subjects (K3R and K8L). Sagittal-plane angles (hip flexion, knee, ankle) reproduced the waveform shape with a small off-set, whereas hip ad/abduction and internal/external rotation showed the largest discrepancies, consistent with previous IMU validation studies (Al Borno et al., 2022; Fan et al., 2022; Jiang et al., 2022). The hip offsets likely resulted from residual sensor-to-segment misalignment during static calibration, propagated by IMUPlacer into the proximal joint angles (Al Borno et al., 2022; Cappozzo et al., 1995).

**Figure 2:**
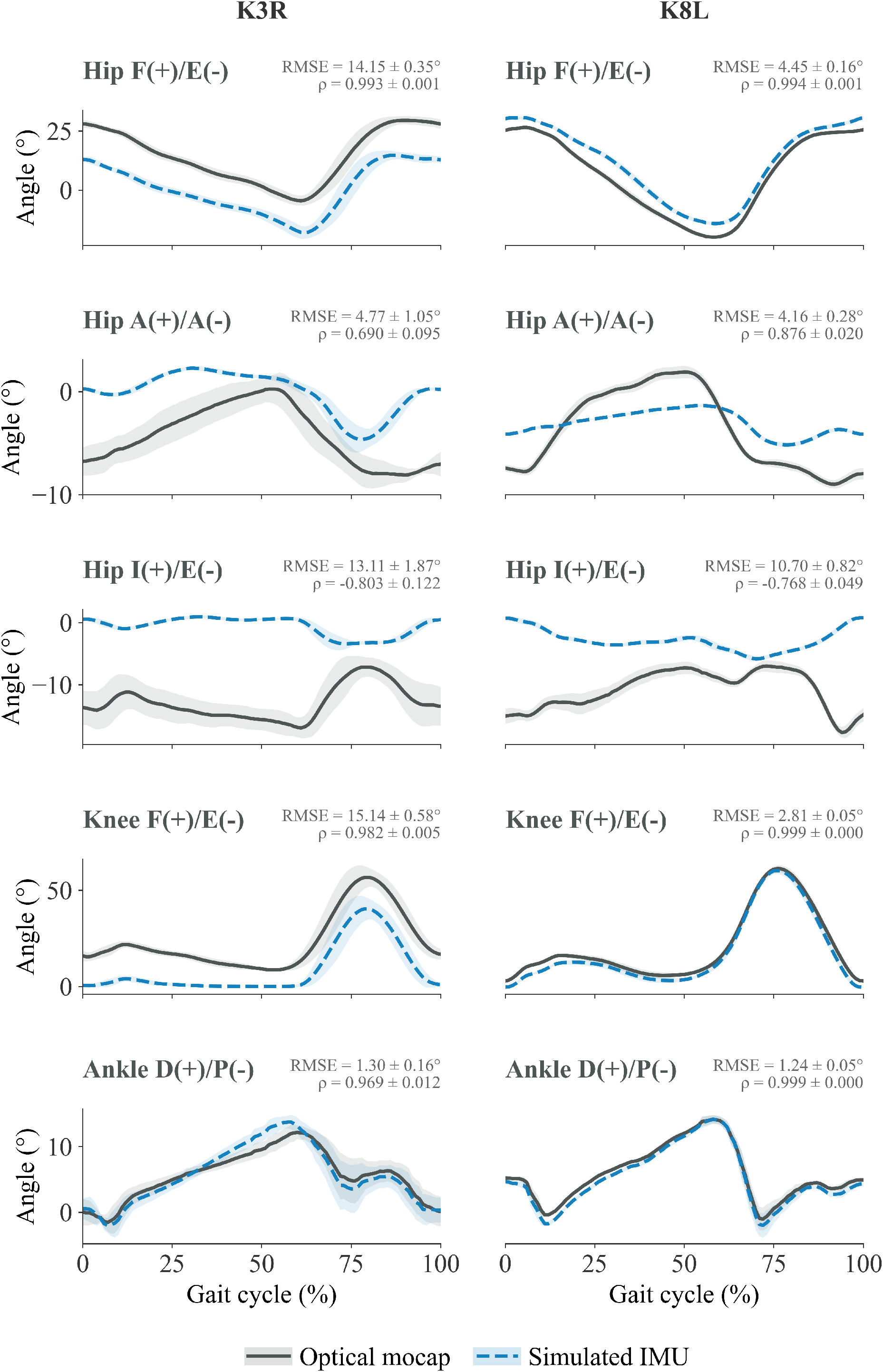
Kinematics comparison for the worst-(K3R) and best-reconstructed (K8L) subjects for level walking over the gait cycle. Solid black: optical mocap reference; dashed blue: SiMPLeSENSE reconstruction. Sagittal angles are reproduced closely (with a small offset), whereas the off-axis hip degrees of freedom show larger deviations. Shaded bands denote ± 1 SD across trials.

Across level walking (LW) and stair descent (SD), all four estimation pipelines reproduced the overall shape and double-peak pattern of the *in vivo* tibiofemoral contact force (Figs. 3, 4). The simulated-IMU estimates closely matched the optical mocap-driven estimates (RMSE_IMU vs optical_ = 0.36 ± 0.22 BW, *ρ* = 0.97 ± 0.03), indicating that replacing optical with inertial inputs did not introduce systematic changes in waveform shape. Compared with the optical pipelines, the IMU-driven methods showed greater trial-to-trial variability, reflecting uncertainty introduced by mesh-based orientation reconstruction. The largest discrepancies occurred at the first force peak in a few subjects (e.g., K3R and K5R), where the IMU-based methods overestimated the *in vivo* force. These were also the subjects with relatively low *in vivo* loads and a stiffer, less regular gait (as reflected by the larger kinematic standard deviations for K3R in Fig. 2).

**Figure 3:**
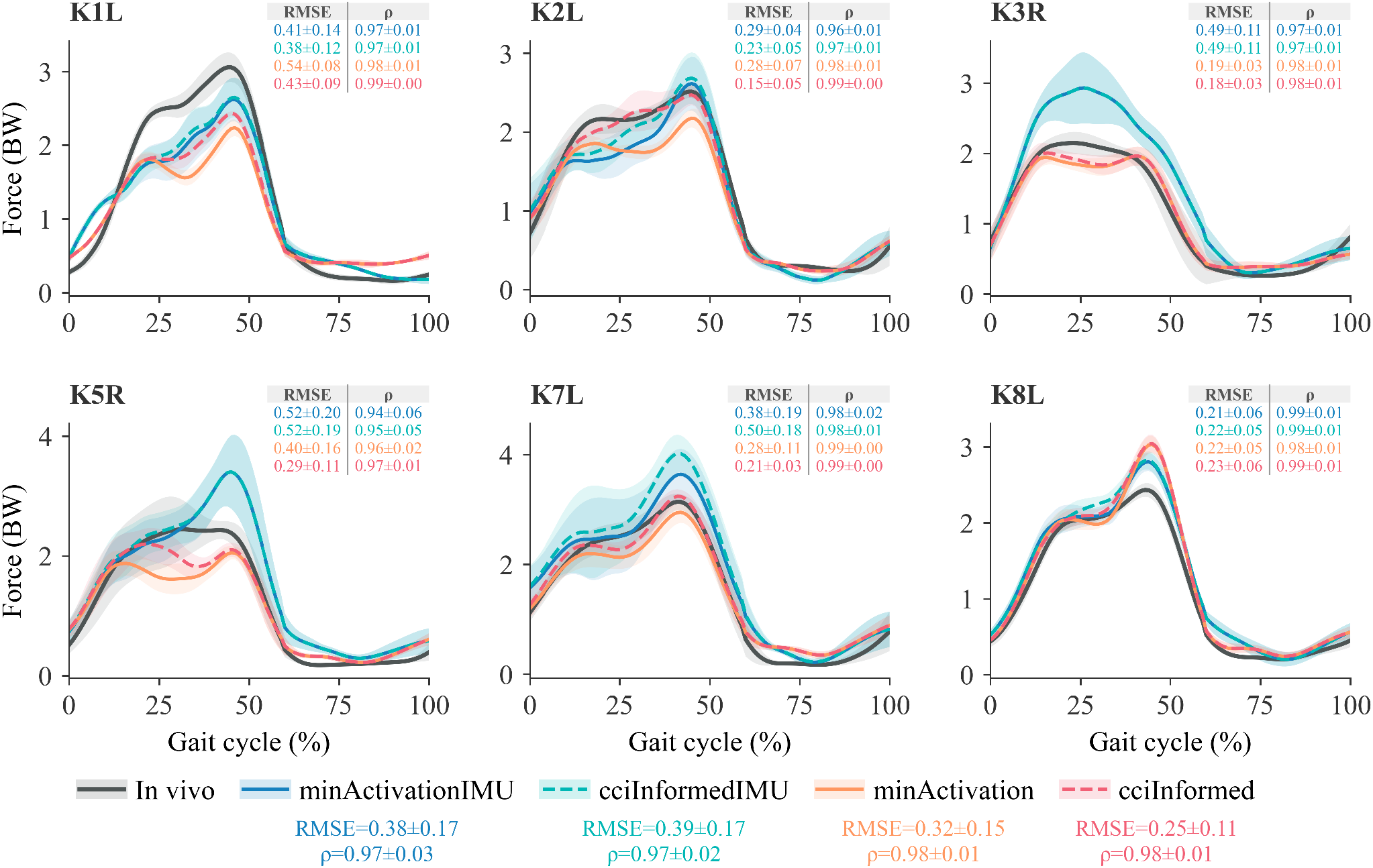
Knee joint compressive forces during level walking for all subjects (K1L–K8L), in bodyweight (BW) over the gait cycle. IMU-driven estimates (*minActivationIMU, cciInformedIMU* ) are compared with the optical mocap-driven estimates of Hörmann et al. (*min-Activation, cciInformed* ) and the *in vivo* measurements. Bands denote ± 1 SD across trials. RMSE and *ρ* (mean ± SD) are reported per-subject within each subplots and overall below figure legend.

**Figure 4:**
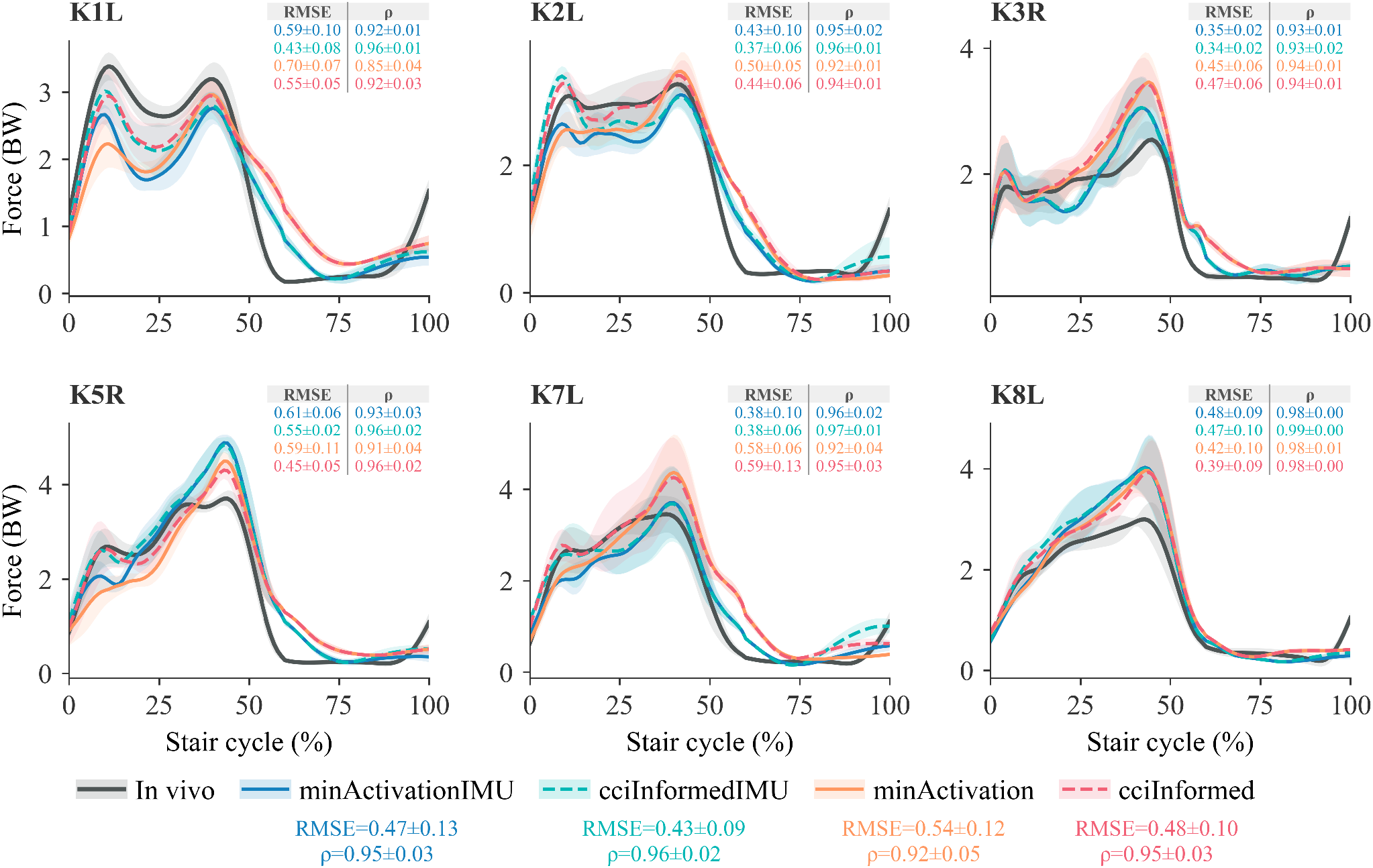
Knee joint compressive forces during stair descent for all subjects (K1L–K8L), in bodyweight (BW) over the gait cycle. IMU-driven estimates (*minActivationIMU, cciInformedIMU* ) are compared with the optical mocap-driven estimates of Hörmann et al. (*min-Activation, cciInformed* ) and the *in vivo* measurements. Bands denote ± 1 SD across trials. RMSE and *ρ* (mean ± SD) are reported per-subject within each subplots and overall below figure legend.

Quantitatively (Fig. 5), agreement was higher for LW than SD. Median RMSE remained below approximately 0.35 BW for LW and increased to roughly 0.4–0.5 BW for SD, while *ρ* exceeded 0.97 for LW and decreased to about 0.92–0.96 for SD. Among the four methods, the optical CCI-informed condition (*cci-Informed* ) achieved the lowest RMSE for LW, whereas the two IMU-based conditions reached comparable correlations despite slightly higher errors.

**Figure 5:**
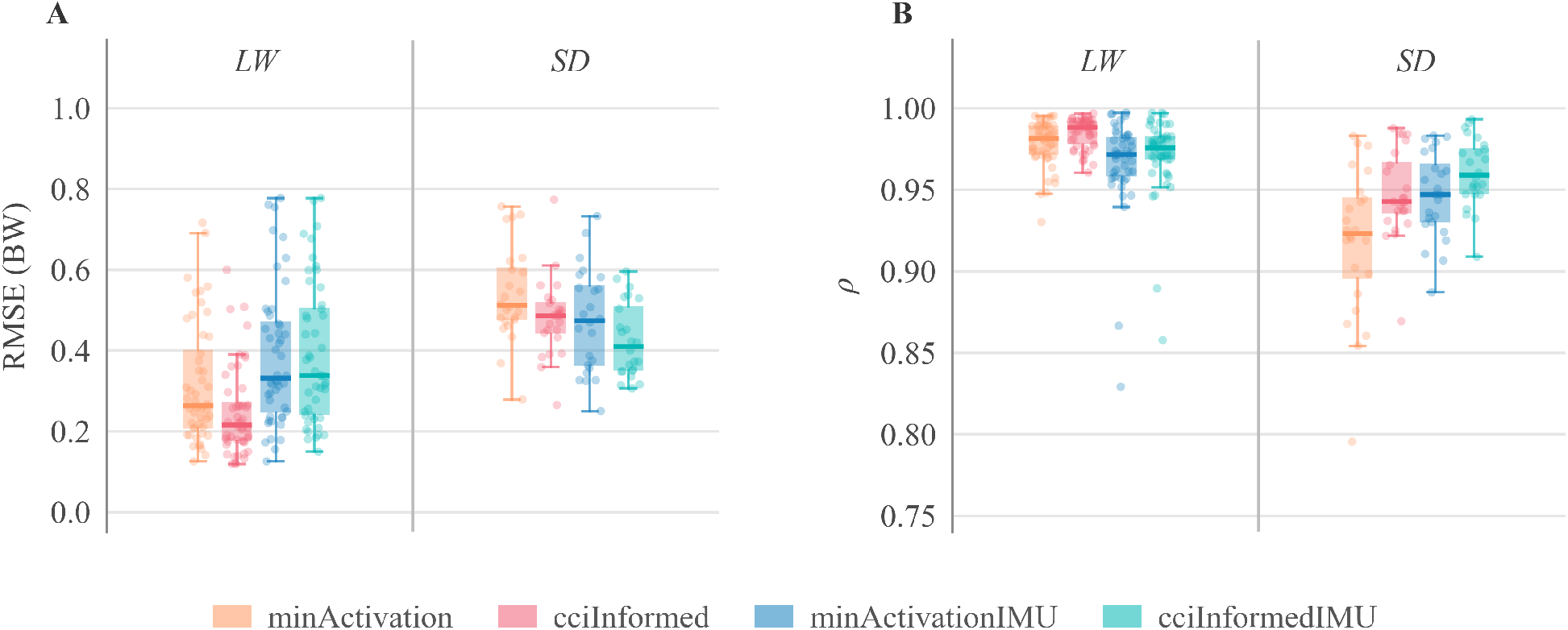
Agreement of IMU-driven estimates (*minActivationIMU, cciInformedIMU* ) and mocap-driven estimates of Hörmann et al. (*minActivation, cciInformed* ) with the *in vivo* measurement, averaged across subjects, for level walking (LW) and stair descent (SD): (A) RMSE in body weight (BW) and (B) Pearson correlation coefficient *ρ*. Boxes show median and interquartile range; dots are individual trials.

Adding realistic IMU orientation noise progressively increased estimation error (Figs. 6, 7). For this analysis, we used *minActivationIMU* as the baseline because it provides the direct IMU counterpart of the conventional *minActivation* method, allowing the effect of orientation error to be assessed independently of the additional CCI constraint. The baseline (*minActivationIMU* ) and single-error (*minActivationIMUError* ) conditions remained close to the *in vivo* curves, whereas doubling the error (*minActivationIMUErrorx2* ) produced clear overestimation of peak force, particularly during stair descent and in subjects with elevated forces (e.g., K3R and K7L). Subject-level RMSE increased consistently across the three conditions (Fig. 8). However, the added error did not affect all subjects equally: it reduced RMSE when baseline estimates underestimated the *in vivo* force (e.g., K1L, K2L and K5R during LW) but substantially increased error when baseline estimates already overpredicted the measured force (most clearly K3R).

**Figure 6:**
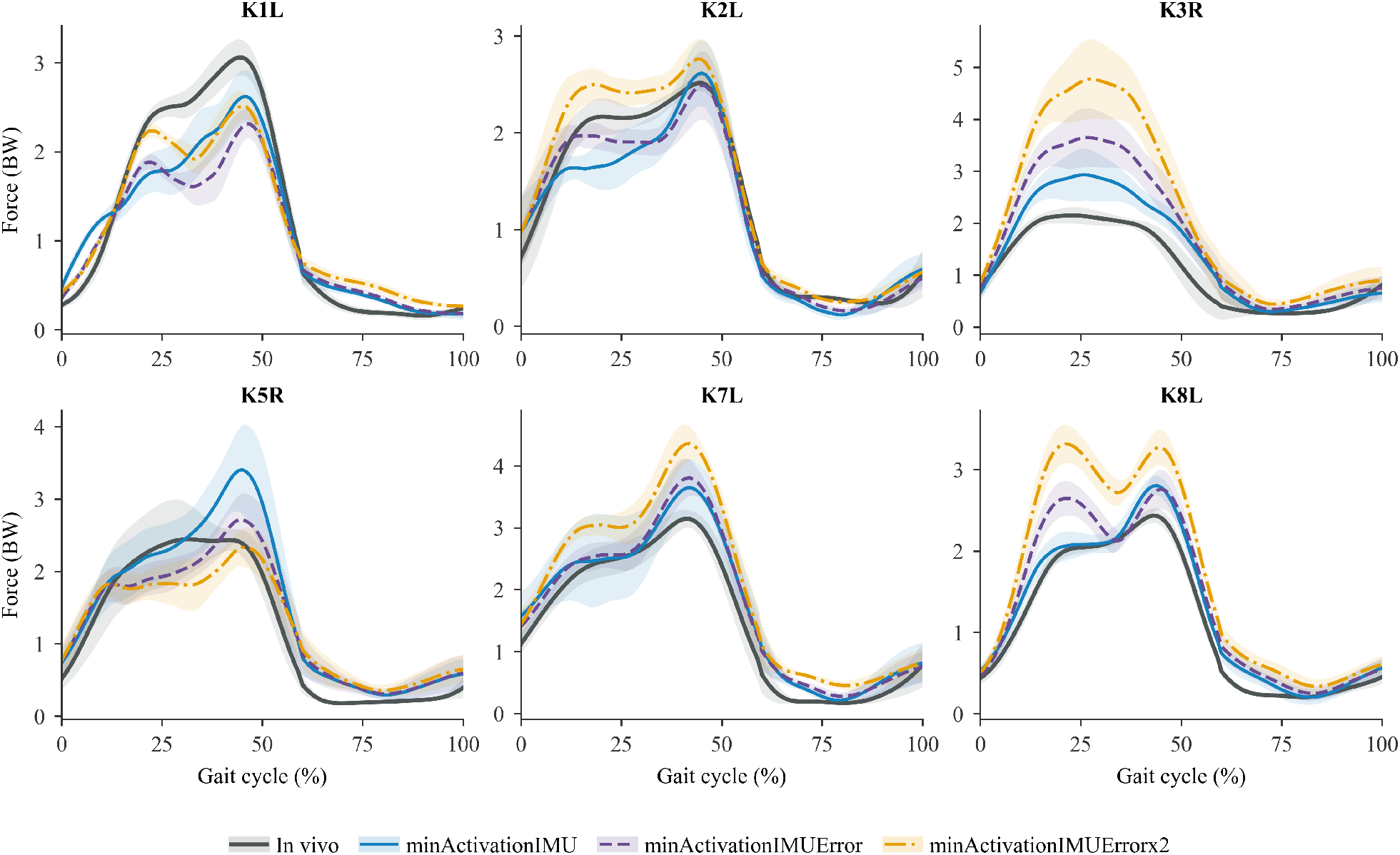
Effect of simulated IMU orientation error on level walking compressive force estimates. The baseline (*minActivationIMU* ) is compared with the addition of Gaussian thigh and shank error (*minActivationIMUError* ; mean and SD from Fan et al. (2022)) and its doubling (*minActivationIMUErrorx2* ), alongside the *in vivo* measurement.

**Figure 7:**
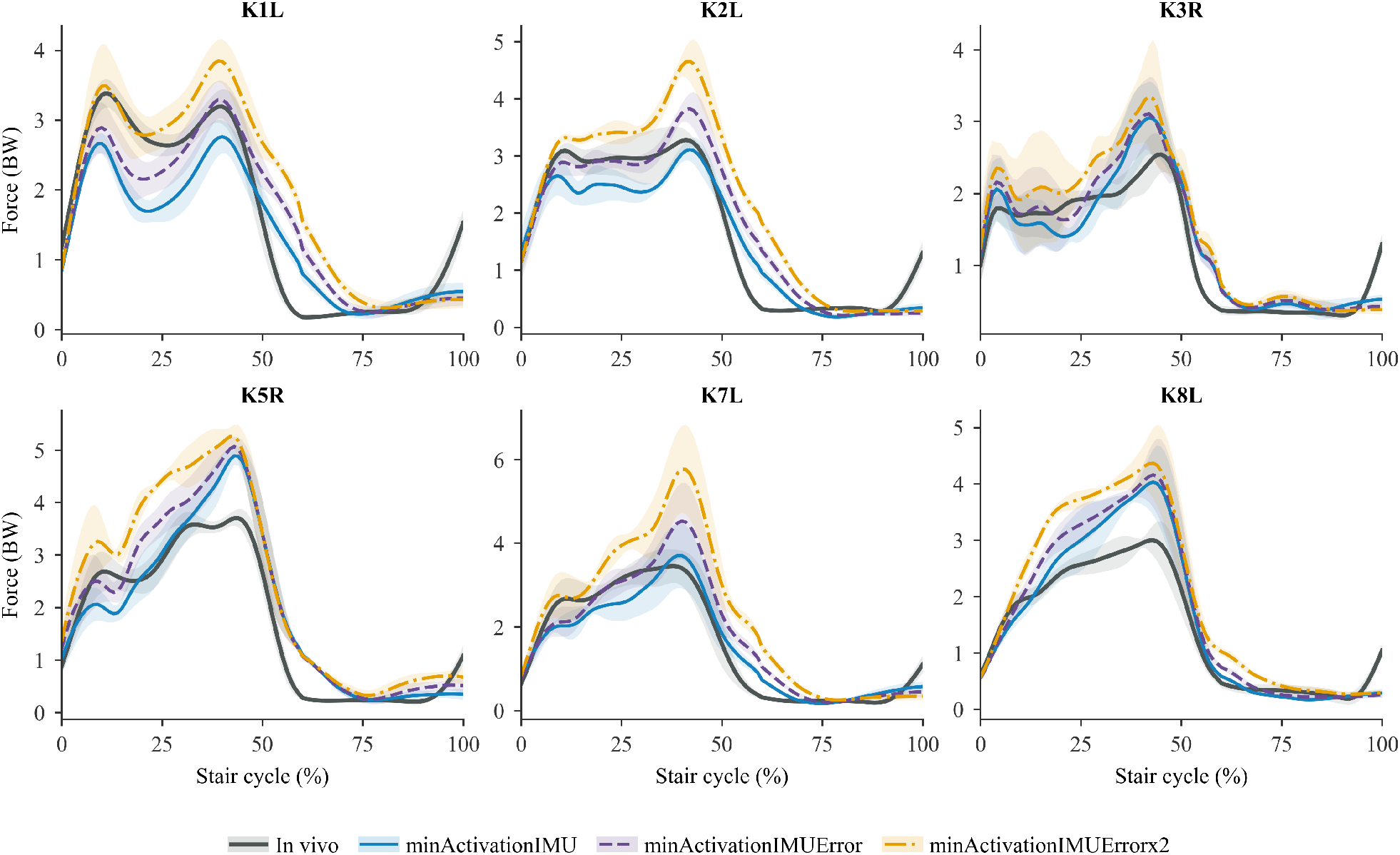
Effect of simulated IMU orientation error on stair descent compressive force estimates. The baseline (*minActivationIMU* ) is compared with the addition of Gaussian thigh and shank error (*minActivationIMUError* ; mean and SD from Fan et al. (2022)) and its doubling (*minActivationIMUErrorx2* ), alongside the *in vivo* measurement.

**Figure 8:**
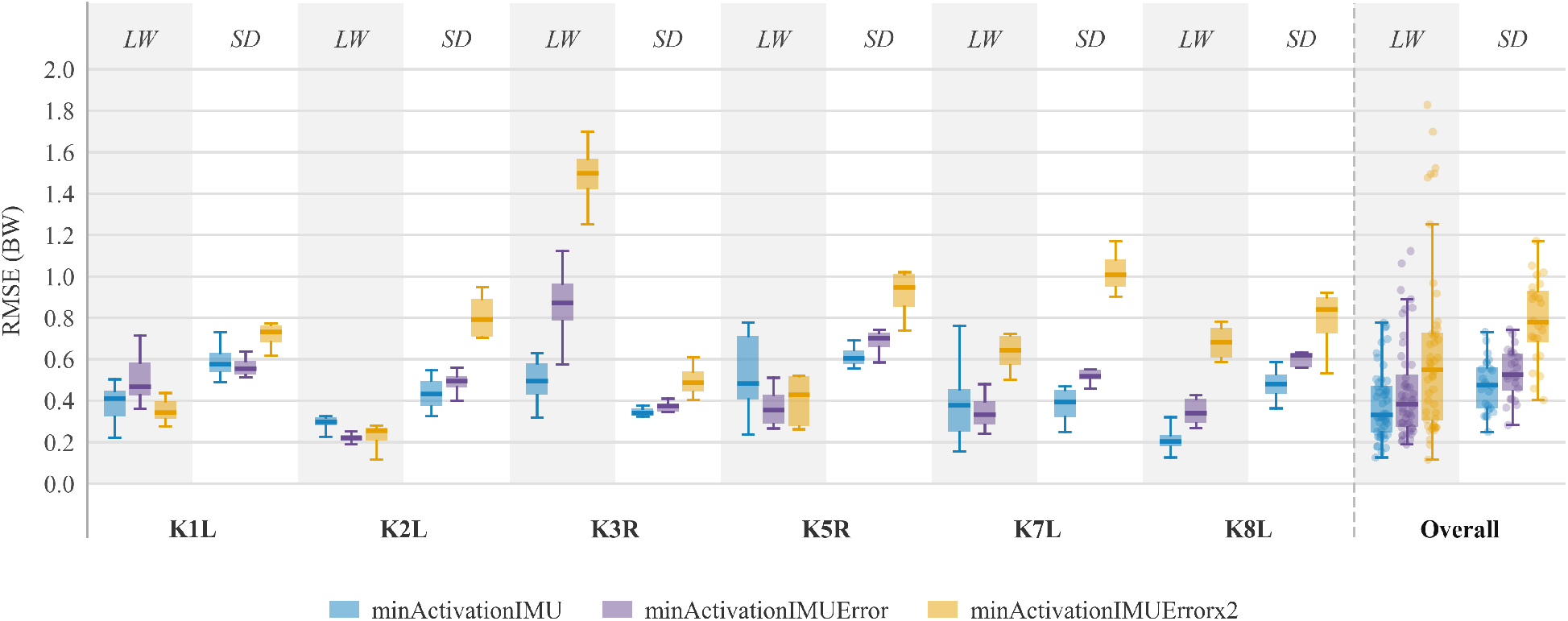
RMSE (BW) of the three noise conditions for each subject and pooled across subjects (Overall), separated by activity, level walking (LW), and stair descent (SD).

## Discussion

This study evaluated whether KCF can be estimated from simulated inertial kinematics with accuracy comparable to an optical, EMG-informed reference, and how robust that estimate remains once realistic sensor-placement errors are introduced. The analysis addressed two questions: how faithfully the SiM-PLeSENSE pipeline reproduces optical mocap-driven KCF, and how the full estimation chain degrades when thigh-and shank-orientation errors are added to the virtual IMUs.

The kinematic reconstruction (Fig. 2) behaved as expected from the IMU literature, with sagittal-plane angles recovered well and the other hip degrees of freedom least accurate (Al Borno et al., 2022; Fan et al., 2022; Jiang et al., 2022). This error distribution is not problematic for contact-force estimation: tibiofemoral loading is governed largely by sagittal-plane knee mechanics, so the degrees of freedom recovered most reliably are also those that matter most for KCF, whereas the larger frontal-and transverse-plane errors, concentrated at the hip, have limited influence on the estimated knee force. The residual sagittal offset, being largely constant, shifts the waveform without distorting its shape, consistent with the high correlations reported below. Because the reconstruction errors were nonetheless somewhat larger than those typically reported for dedicated physical IMU systems, the downstream KCF results are a conservative test of the method.

The first part of the analysis showed good agreement between the optical mocap-driven and simulated-IMU-driven KCF. The IMU-driven waveforms tracked the optical estimates closely (RMSE_IMU vs optical_ = 0.36 ± 0.22 BW, *ρ* = 0.97 ± 0.03), with agreement that was, higher for stair descent than for level walking. This is a central result: despite the kinematic discrepancies, the estimated KCF changed less than might be expected, suggesting that, within the RMR pipeline, joint loading is influenced more strongly by the GRF than by the kinematic differences introduced by the inertial reconstruction. The simulated-IMU estimates did show greater trial-to-trial variability: the virtual IMUs attach to different mesh faces of the SMPL surface, so each segment frame samples a slightly different realization of soft-tissue motion, analogous to the soft-tissue artefact that dominates skin-marker-based kinematics (Leardini et al., 2004; Cappozzo et al., 1995). Peak force was overestimated in K3R and K5R, both of whom walked with a stiffer, more irregular pattern, reflected in the greater kinematic variability of K3R in Fig. 2. In such gait, small kinematic changes can have a disproportional effect on predicted loading, and the effect is amplified for K3R, whose low *in vivo* KCF turns a given absolute error into a large relative error.

The two solver variants differed mainly by activity. The CCI-informed condition (*cciInformed* ) performed almost identically to the minimum-activation condition (*minActivation*) for level walking and improved on it only slightly for stair descent. This is consistent with the role of co-contraction in knee loading: antagonist co-activation contributes more during the more demanding stairdescent task, so constraining the solution toward the measured co-contraction has a larger effect there, whereas level walking is already well captured by the minimum-activation objective and leaves little room for improvement (Belli et al., 2023; Hörmann et al., 2026). The same ordering held for the IMU-driven estimates, indicating that the benefit of the CCI term is preserved when the input kinematics come from simulated inertial data rather than optical motion capture. OA patients typically exhibit elevated co-contraction compared to healthy controls, though this is normalized following total knee arthroplasty (Fallah-Yakhdani et al., 2012). Given the role of excessive co-contraction in OA progression, the present method could be particularly valuable for monitoring joint loading in pre-arthroplasty OA populations.

In the second part of the analysis we superimposed orientation errors on the thigh and shank IMUs to mimic the misalignment and orientation errors of a physical deployment. Under the realistic magnitudes reported by Fan et al. (2022), the pipeline still performed well, with overall median RMSE rising only modestly (median difference of 0.065 BW for level walking and 0.059 BW for stair descent) from baseline (Fig. 8). For some subjects the added error improved agreement, because their baseline had underpredicted the *in vivo* force, whereas subjects whose baseline already overpredicted (most clearly K3R) deteriorated. Doubling the error revealed the method’s limits: peak forces were systematically overestimated and trial-to-trial variability widened. Even then, the doubled error reduced RMSE for a subset of subjects (K1L, K2L, and K5R in LW), again because it pushed previously underpredicted estimates toward the measured values. Overall, increasing sensor orientation error tended to increase predicted force and its variability, while the effect on RMSE depended on each subject’s baseline bias.

Studies that drive musculoskeletal models from inertial inputs report errors in a similar range, although most compare against optical estimates rather than *in vivo* forces. Konrath et al. (2019) estimated tibiofemoral contact force from inertial motion capture with RMSE of 0.4 to 1.0 BW and moderate-to-strong correlations against an optical-driven model, and Di Raimondo et al. (2023) reported IMU-based peak forces that did not differ significantly from motion-capture estimates (RMSE 0.15 - 0.40 BW). Our IMU-to-optical agreement (RMSE_IMU vs optical_ = 0.36 ± 0.22 BW, *ρ* = 0.97 ± 0.03) falls within this range, indicating that the inertial reconstruction adds error comparable to existing IMU pipelines. The methodological distinction is that, by generating the virtual sensors directly on the body mesh, the sensor-free SiMPLeSENSE design isolates the contribution of the inertial reconstruction itself and allows comparison against *in vivo* rather than optical references, while the perturbation analysis quantifies how the estimate degrades as orientation error grows, complementing analytical sensitivity studies of IMU knee kinematics (Fan et al., 2022; Jiang et al., 2022).

Although the present findings are promising, several limitations should be acknowledged. First, the study was conducted on a relatively small cohort, all of whom had undergone total knee replacement, which may limit the generalizability of the results to broader or populations showing more co-contraction.

The cohort was also predominantly male (five of six participants), so any sexrelated differences in knee loading could not be assessed and the findings should be interpreted with this imbalance in mind (van der Kruk, 2025). Second, the analysis did not rely on signals acquired from physical IMUs. Instead, IMU orientations were simulated, which may not fully capture the noise, drift, and measurement variability inherent to real-world sensor recordings. Future work should therefore aim to validate the proposed approach using in-vivo joint force measurements and signals collected from actual IMU sensors, as well as to confirm its robustness in a larger and more heterogeneous cohort.

Observing overall RMSE and *ρ* outcomes, the RMR solver proved robust to changes in the input kinematics, including the relatively large and even doubled errors imposed here, yielding KCF estimates close to the optical motion capture gold standard. The observed KCF errors remain below the minimum detectable change threshold of 0.66 BW reported for EMG-driven knee contact force models in healthy populations (Gardinier et al., 2013), suggesting that the differences may not reflect meaningful changes in joint loading. This indicates that the estimation accuracy is within a clinically interpretable range. Future work should determine whether this accuracy translates to reliable detection of meaningful changes in joint loading, including sensitivity and specificity for load-change detection. With a modest loss in accuracy, KCF could be estimated using a fully portable setup combining inertial motion capture, limited EMG channels to capture muscle co-contraction, and ground reaction forces. Replacing force plates with instrumented force shoes (Schepers et al., 2007) would enable laboratory-independent monitoring of joint loading, supporting longitudinal assessment of rehabilitation outcomes and cumulative loading in patients at risk of osteoarthritis progression.

## Acknowledgements

This publication is part of the project LoaD (projectnr. NWA1389.20.009) of the NWA-ORC research programme which is420 (partly) financed by the Dutch Research Council (NWO).

## Declaration of Generative AI and AI-assisted Technologies in the Manuscript Preparation Process

During the preparation of this work, the authors used ChatGPT for language editing and refinement of the manuscript. The authors reviewed and edited the output as needed and take full responsibility for the content of the published article.

## References

Al Borno, M., O’Day, J., Ibarra, V., Dunne, J., Seth, A., Habib, A., Ong, C., Hicks, J., Uhlrich, S., & Delp, S. (2022). OpenSense: An open-source toolbox for inertial-measurement-unit-based measurement of lower extremity kinematics over long durations. Journal of NeuroEngineering and Rehabilitation, 19 (1), 22. 10.1186/s12984-022-01001-x

Andriacchi, T. P., & Mündermann, A. (2006). The role of ambulatory mechanics in the initiation and progression of knee osteoarthritis. Current Opinion in Rheumatology, 18, 514–518.

Belli, I., Joshi, S., Prendergast, J. M., Beck, I., Della Santina, C., Peternel, L., & Seth, A. (2023). Does enforcing glenohumeral joint stability matter? A new rapid muscle redundancy solver highlights the importance of non-superficial shoulder muscles. PLOS ONE, 18 (11), Article e0295003. 10.1371/journal.pone.0295003

Cappozzo, A., Catani, F., Della Croce, U., & Leardini, A. (1995). Position and orientation in space of bones during movement: Anatomical frame definition and determination. Clinical Biomechanics, 10 (4), 171–178. 10.1016/0268-0033(95)91394-T

Crowninshield, R. D., & Brand, R. A. (1981). A physiologically based criterion of muscle force prediction in locomotion. Journal of Biomechanics, 14, 793–801.

Delp, S. L., Anderson, F. C., Arnold, A. S., Loan, P., Habib, A., John, C. T., Guendelman, E., & Thelen, D. G. (2007). OpenSim: Open-source software to create and analyze dynamic simulations of movement. IEEE Transactions on Biomedical Engineering, 54 (11), 1940–1950. 10.1109/tbme.2007.901024

Di Raimondo, G., Willems, M., Killen, B. A., Havashinezhadian, S., Turcot, K., Vanwanseele, B., & Jonkers, I. (2023). Peak tibiofemoral contact forces estimated using IMU-based approaches are not significantly different from motion capture-based estimations in patients with knee osteoarthritis. Sensors, 23 (9), Article 4484. 10.3390/s23094484

Dorschky, E., Nitschke, M., Seifer, A.-K., van den Bogert, A. J., & Eskofier, B. M. (2019). Estimation of gait kinematics and kinetics from inertial sensor data using optimal control of musculoskeletal models. Journal of Biomechanics, 95, Article 109278. 10.1016/j.jbiomech.2019.07.022

Dorschky, E., Nitschke, M., Martindale, C. F., van den Bogert, A. J., Koelewijn, A. D., & Eskofier, B. M. (2020). CNN-based estimation of sagittal plane walking and running biomechanics from measured and simulated inertial sensor data. Frontiers in Bioengineering and Biotechnology, 8, Article 604. 10.3389/fbioe.2020.00604

Fallah-Yakhdani, H. R., Abbasi-Bafghi, H., Meijer, O. G., Bruijn, S. M., van den Dikkenberg, N., Benedetti, M. G., & van Dieën, J. H. (2012). Determinants of co-contraction during walking before and after arthroplasty for knee osteoarthritis. Clinical Biomechanics, 27 (5), 485–494. 10.1016/j.clinbiomech.2011.11.006

Fan, B., Li, Q., Tan, T., Kang, P., & Shull, P. B. (2022). Effects of IMU Sensor-to-Segment Misalignment and Orientation Error on 3-D Knee Joint Angle Estimation. IEEE Sensors Journal, 22 (3), 2543–2552. 10.1109/jsen.2021.3137305

Fregly, B. J., Besier, T. F., Lloyd, D. G., Delp, S. L., Banks, S. A., Pandy, M. G., & D’Lima, D. D. (2012). Grand challenge competition to predict in vivo knee loads. Journal of Orthopaedic Research, 30 (4), 503–513. 10.1002/jor.22023

Gardinier, E. S., Manal, K., Buchanan, T. S., & Snyder-Mackler, L. (2013). Minimum detectable change for knee joint contact force estimates using an EMG-driven model. Gait & Posture, 38 (4), 1051–1053. 10.1016/j.gaitpost.2013.03.014

Hao, Y., Lou, X., Wang, B., & Zheng, R. (2024). CROMOSim: A Deep Learning-Based Cross-Modality Inertial Measurement Simulator. IEEE Transactions on Mobile Computing, 23 (1), 302–312. 10.1109/TMC.2022.3230370

Heinlein, B., Kutzner, I., Graichen, F., Bender, A., Rohlmann, A., Halder, A. M., Beier, A., & Bergmann, G. (2009). ESB Clinical Biomechanics Award 2008: Complete data of total knee replacement loading for level walking and stair climbing measured in vivo with a follow-up of 6–10 months. Clinical Biomechanics, 24 (4), 315–326. 10.1016/j.clinbiomech.2009.01.011

Hoang, H. X., Diamond, L. E., Lloyd, D. G., & Pizzolato, C. (2019). A calibrated EMG-informed neuromusculoskeletal model can appropriately account for muscle co-contraction in the estimation of hip joint contact forces in people with hip osteoarthritis. Journal of Biomechanics, 83, 134–142. 10.1016/j.jbiomech.2018.11.042

Hörmann, S., Tümer, N., Zadpoor, A. A., & Seth, A. (2026). Co-contraction index improves estimates of knee joint contact forces: A musculoskeletal modelling study. bioRxiv. 10.64898/2026.05.28.728089

Imani Nejad, Z., Khalili, K., Hosseini Nasab, S. H., Schütz, P., Damm, P., Trepczynski, A., Taylor, W. R., & Smith, C. R. (2020). The capacity of generic musculoskeletal simulations to predict knee joint loading using the CAMS-Knee datasets. Annals of Biomedical Engineering, 48 (4), 1430–1440. 10.1007/s10439-020-02465-5

Jiang, C., Yang, Y., Mao, H., Yang, D., & Wang, W. (2022). Effects of dynamic IMU-to-segment misalignment error on 3-DOF knee angle estimation in walking and running. Sensors, 22 (22), Article 9009. 10.3390/s22229009

Karatsidis, A., Bellusci, G., Schepers, H. M., de Zee, M., Andersen, M. S., & Veltink, P. H. (2017). Estimation of ground reaction forces and moments during gait using only inertial motion capture. Sensors, 17 (1), Article 75. 10.3390/s17010075

Konrath, J. M., Karatsidis, A., Schepers, H. M., Bellusci, G., de Zee, M., & Andersen, M. S. (2019). Estimation of the knee adduction moment and joint contact force during daily living activities using inertial motion capture. Sensors, 19 (7), Article 1681. 10.3390/s19071681

van der Kruk, E. (2025). BIASMECHANICS: Does an unconscious bias still persist in biomechanics, positioning males as the default in human research? A meta-analysis on the Journal of Biomechanics 2024 publications. Journal of Biomechanics, 181, 112560. 10.1016/j.jbiomech.2025.112560

Kumar, D., Manal, K., & Rudolph, K. S. (2012). Knee joint loading during gait in healthy controls and individuals with knee osteoarthritis. Osteoarthritis and Cartilage, 21 (2), 298–305. 10.1016/j.joca.2012.11.008

Kutzner, I., Heinlein, B., Graichen, F., Bender, A., Rohlmann, A., Halder, A., Beier, A., & Bergmann, G. (2010). Loading of the knee joint during activities of daily living measured in vivo in five subjects. Journal of Biomechanics, 43 (11), 2164–2173. 10.1016/j.jbiomech.2010.03.046

Lai, A. K. M., Arnold, A. S., & Wakeling, J. M. (2017). Why are antagonist muscles co-activated in my simulation? A musculoskeletal model for analysing human locomotor tasks. Annals of Biomedical Engineering, 45, 2762–2774. 10.1007/s10439-017-1920-7

Laidig, D., & Seel, T. (2023). VQF: Highly accurate IMU orientation estimation with bias estimation and magnetic disturbance rejection. Information Fusion, 91, 187–204. 10.1016/j.inffus.2022.10.014

Leardini, A., Chiari, L., Della Croce, U., & Cappozzo, A. (2004). Human movement analysis using stereophotogrammetry. Part 3: Soft tissue artifact assessment and compensation. Gait & Posture, 21 (2), 212–225. 10.1016/j.gaitpost.2004.05.002

Lloyd, D. G., & Besier, T. F. (2003). An EMG-driven musculoskeletal model to estimate muscle forces and knee joint moments in vivo. Journal of Biomechanics, 36, 765–776.

Loper, M., Mahmood, N., Romero, J., Pons-Moll, G., & Black, M. J. (2023). SMPL: A skinned multi-person linear model. Seminal Graphics Papers: Pushing the Boundaries, 2. 10.1145/3596711.3596800

Mahmood, N., Ghorbani, N., Troje, N. F., Pons-Moll, G., & Black, M. J. (2019). AMASS: Archive of Motion Capture As Surface Shapes. In Proceedings of the IEEE/CVF International Conference on Computer Vision (ICCV) (pp. 5442–5451). 10.1109/ICCV.2019.00554

Manal, K., & Buchanan, T. S. (2013). An electromyogram-driven musculoskeletal model of the knee to predict in vivo joint contact forces during normal and novel gait patterns. Journal of Biomechanical Engineering, 135 (2), Article 021014. 10.1115/1.4023457

Merletti, R. (1999). Standards for reporting EMG data. Journal of Electromyography and Kinesiology, 9 (1), III–IV.

Mundt, M., Koeppe, A., David, S., Witter, T., Bamer, F., Potthast, W., & Markert, B. (2020). Estimation of gait mechanics based on simulated and measured IMU data using an artificial neural network. Frontiers in Bioengineering and Biotechnology, 8, Article 41. 10.3389/fbioe.2020.00041

Pelegrinelli, A. R. M., Catelli, D. S., Kowalski, E., Lamontagne, M., & Moura, F. A. (2023). Comparing three generic musculoskeletal models to estimate the tibiofemoral reaction forces during gait and sit-to-stand tasks. Medical Engineering & Physics, 122, Article 104074. 10.1016/j.medengphy.2023.104074

Picerno, P. (2017). 25 years of lower limb joint kinematics by using inertial and magnetic sensors. Gait & Posture, 51, 239–246. 10.1016/j.gaitpost.2016.11.008

Pizzolato, C., Lloyd, D. G., Sartori, M., Ceseracciu, E., Besier, T. F., Fregly, B. J., & Reggiani, M. (2015). CEINMS: A toolbox to investigate the influence of different neural control solutions on the prediction of muscle excitation and joint moments during dynamic motor tasks. Journal of Biomechanics, 48 (14), 3929–3936. 10.1016/j.jbiomech.2015.09.021

Princelle, D., Viceconti, M., & Davico, G. (2025). EMG-Informed Neuromusculoskeletal Simulations Increase the Accuracy of the Estimation of Knee Joint Contact Forces During Sub-optimal Level Walking. Annals of Biomedical Engineering. 10.1007/s10439-025-03713-2

Rajagopal, A., Dembia, C. L., DeMers, M. S., Delp, D. D., Hicks, J. L., & Delp, S. L. (2016). Full-Body Musculoskeletal Model for Muscle-Driven Simulation of Human Gait. IEEE Transactions on Biomedical Engineering, 63 (10), 2068–2079. 10.1109/TBME.2016.2586891

Rudolph, K. S., Axe, M. J., Buchanan, T. S., Scholz, J. P., & Snyder-Mackler, L. (2001). Dynamic stability in the anterior cruciate ligament deficient knee. Knee Surgery, Sports Traumatology, Arthroscopy, 8, 62–71. 10.1007/s001670000166

Sartori, M., Reggiani, M., Farina, D., & Lloyd, D. G. (2012). EMG-driven forward-dynamic estimation of muscle force and joint moment about multiple degrees of freedom in the human lower extremity. PLOS ONE, 7 (12), Article e52618. 10.1371/journal.pone.0052618

Schepers, H. M., Koopman, H. F. J. M., & Veltink, P. H. (2007). Ambulatory Assessment of Ankle and Foot Dynamics. IEEE Transactions on Biomedical Engineering, 54 (5), 895–902. 10.1109/TBME.2006.889769

Schepers, M., Giuberti, M., & Bellusci, G. (2018). Xsens MVN: Consistent tracking of human motion using inertial sensing (Tech. Rep. No. MV0424P.A). Xsens Technologies.

Seth, A., Hicks, J. L., Uchida, T. K., Habib, A., Dembia, C. L., Dunne, J. J., Ong, C. F., DeMers, M. S., Rajagopal, A., Millard, M., Hamner, S. R., Arnold, E. M., Yong, J. R., Lakshmikanth, S. K., Sherman, M. A., Ku, J. P., & Delp, S. L. (2018). OpenSim: Simulating musculoskeletal dynamics and neuromuscular control to study human and animal movement. PLOS Computational Biology, 14 (7), Article e1006223. 10.1371/journal.pcbi.1006223

Shuster, M. D. (1993). A survey of attitude representations. Journal of the Astronautical Sciences, 41, 439–517.

Taylor, W. R., Schütz, P., Bergmann, G., List, R., Postolka, B., Hitz, M., Dymke, J., Damm, P., Duda, G., Gerber, H., Schwachmeyer, V., Hosseini Nasab, S. H., Trepczynski, A., & Kutzner, I. (2017). A comprehensive assessment of the musculoskeletal system: The CAMS-Knee dataset. Journal of Biomechanics, 65, 32–39. 10.1016/j.jbiomech.2017.09.022

Uhlrich, S. D., Mazzoli, V., Silder, A., Finlay, A. K., Kogan, F., Gold, G. E., Delp, S. L., Beaupre, G. S., & Kolesar, J. A. (2025). Personalised gait retraining for medial compartment knee osteoarthritis: a randomised controlled trial. The Lancet Rheumatology, 7 (10), e708–e718. 10.1016/S2665-9913(25)00151-1

Uhlrich, S. D., Jackson, R. W., Seth, A., Kolesar, J. A., & Delp, S. L. (2022). Muscle coordination retraining inspired by musculoskeletal simulations reduces knee contact force. Scientific Reports, 12, 9842. 10.1038/s41598-022-13386-9

